## Supplementary figures and tables for "Metabotropic signaling within somatostatin interneurons controls transient thalamocortical inputs during development"

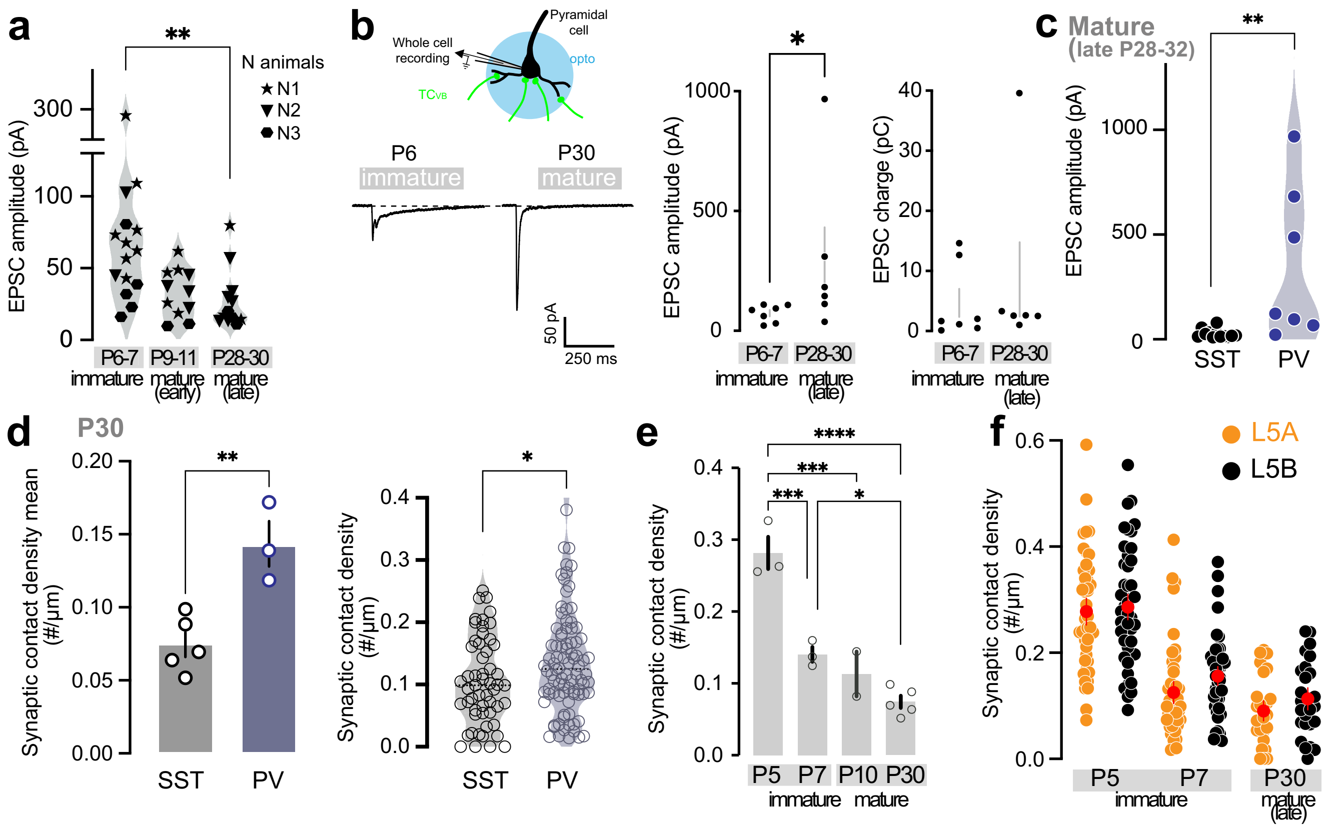


### Supplementary Fig.1: TC circuit wiring onto SST and PV cINs

**a**, EPSC amplitudes from Fig. 1c labeled with biological N (n cells recorded per animal). **b**, Example response traces of EPSCs recorded from excitatory pyramidal neuron in L5, as control for SST cIN recordings in the same slice. EPSC amplitude is increased during development, as previously described. Mann-Whitney test (p-value=0.022). **c**, EPSC amplitudes from SST and PV cINs during late mature stage (P28-32) upon optogenetic stimulation of $TC_{VB}$.fibers. As expected, $TC_{VB}$ are strongly connected to PV cINs compared to SST cINs at maturation. Mann-Whitney test (p-value=0.0012). **d**, TC synaptic density (VGluT2+/Homer1+) reflects the physiological strength of TC inputs to SST and PV cINs. Left: Average per animal of TC synaptic contact density. Student t-test (p-value=0.005; SST 0.074±0.008 N=5; PV 0.143±0.015). Right: synaptic contact density per cell. Student t-test (p-value=0.0147; SST:0.105±0.009 n=54 N=5 PV: 0.140±0.009 n=90 N=3).**e**, Average per animal of TC synaptic contact density (VGluT2+/Homer1+) onto SST cINs showing a decrease of synapse numbers during development. One-way ANOVA (p-value <0.0001) followed by Tukey's multiple comparison tests (p-values for P5-P7=0.0008, P5-P30<0.0001, P7-P30=0.0425, P5-10=0.0005, P7-P10=0.715, P10-P30=0.4001; P5: 0.282±0.023 N=3, P7: 0.141±0.010 N=3, P10: 0.144±0.015 N=2, P30: 0.074±0.008 N=5). **f**, The same quantification was performed based on the localization of the SST cINs in L5A vs L5B. No significant difference in TC synaptic contact number was detected between the layers at each age.

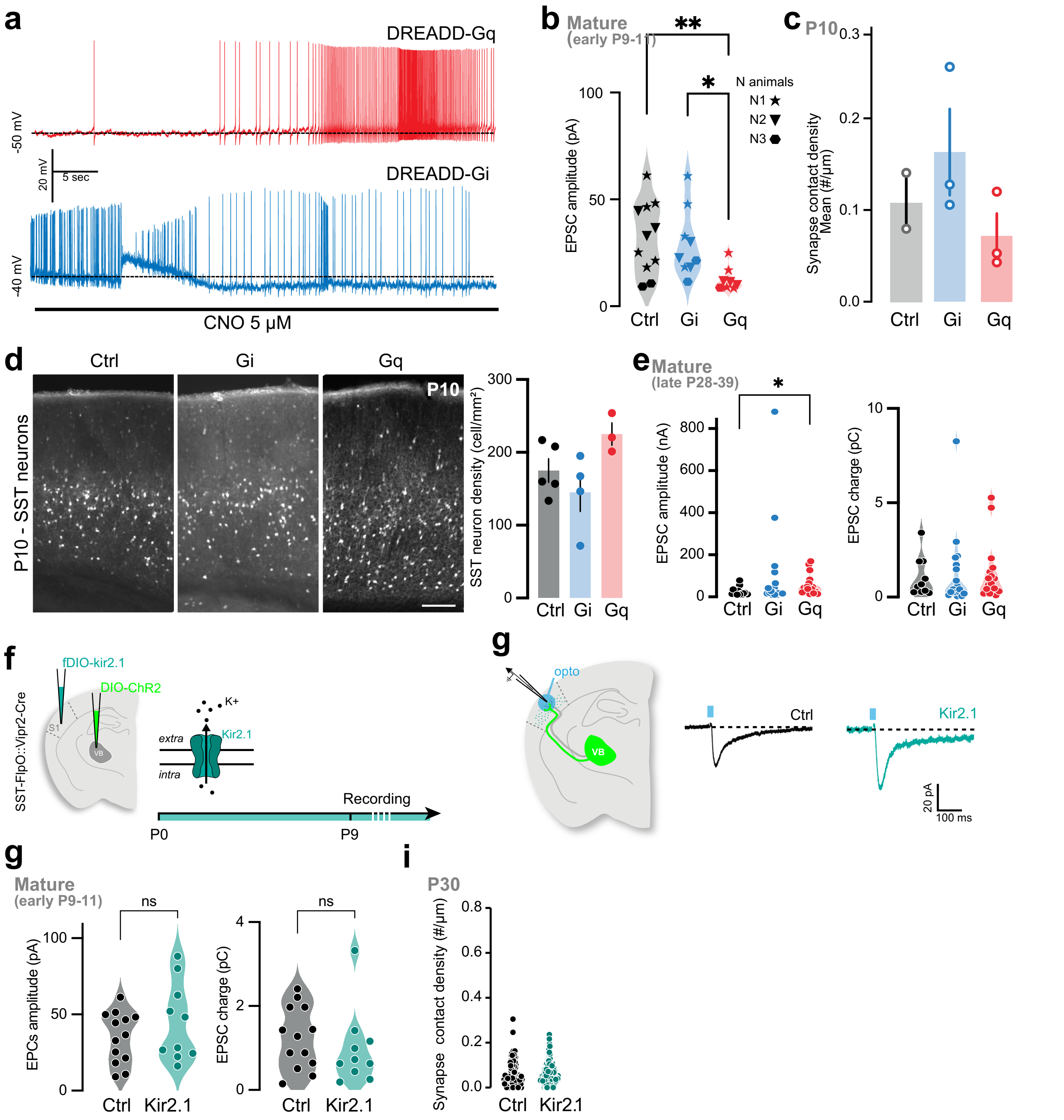

**Supplementary Fig. 2: TC circuit wiring upon chemogenetic manipulation of SST cINs**

**a**, Example of depolarization and increase in firing of DREADD-Gq(+) SST cINs and hyperpolarization and decrease in firing of DREADD-Gi(+) SST cINs upon bath application of CNO, on P10 slices from chronically activated brains. **b**, EPSC amplitudes from Fig. 2d, labeled for biological N (n cells recorded per animal). **c**, Average per animal of synaptic contact density onto SST cINs (Ctrl from Fig.1: 0.113±0.031 N=2, Gi: 0.163±0.47 N=3; Gq: 0.072±0.024 N=3). **d**, S1 SST cINs density at P10 after chronic activation of Dreadd-Gi/Gq (P1-8) and Ctrl (vehicle). Scale bar = 100 µm. Cell number/mm$^2$ of Ctrl (Gi(+)-saline), Gi(+) and Gq(+) SST cINs. Non-significant One-way ANOVA (Ctrl: 173±11.6 N=6; Gi: 184.5±31 N=5; Gq: 225±12.5 N=3). **e**, EPSC peak amplitude and charges of Ctrl, Gi- and Gq-expressing SST cINs in response to VB fiber stimulations at P30. Kruskal-Wallis (p-value=0.0435) followed by Dunn's multiple comparisons test (p-value for Ctrl-Gq=0.038; Ctrl: 25.34±5.04 n=15 N; Gi: 98.42±47.71 n=19; Gq: 57.38±11.75 n=17). **f**, Strategy for expression of potassium ionotropic channel Kir2.1 in SST cINs. AAV-driven Cre-dependent ChR2 expression in VB is controlled by Vipr2-Cre mouse line. Flp-dependent Kir2.1 (fDIO) expression in SST cINs is controlled by SST-FlpO mouse line. **g**, Recording of SST Kir2.1(+) cINs upon optogenetic stimulation of VB fibers. Averaged response trace of example neurons from Ctrl and Kir2.1(+) cINs. EPSC peak amplitudes and charges between control (from Fig. 2) and Kir2.1 expressing SST cINs in L5 at P10 do not show significant change. Mann-Whitney test (Amplitudes: Ctrl 35.08±4.67; Kir2.1 44.88±8.07; Charges: Ctrl 0.97±0.29; .24±0.21). **i**, Synaptic contact density onto SST cINs at P30 (Ctrl: 0.062±0.0005 n=195 N=3; Kir2,1: 0.079±0.003 n=230 N=6).

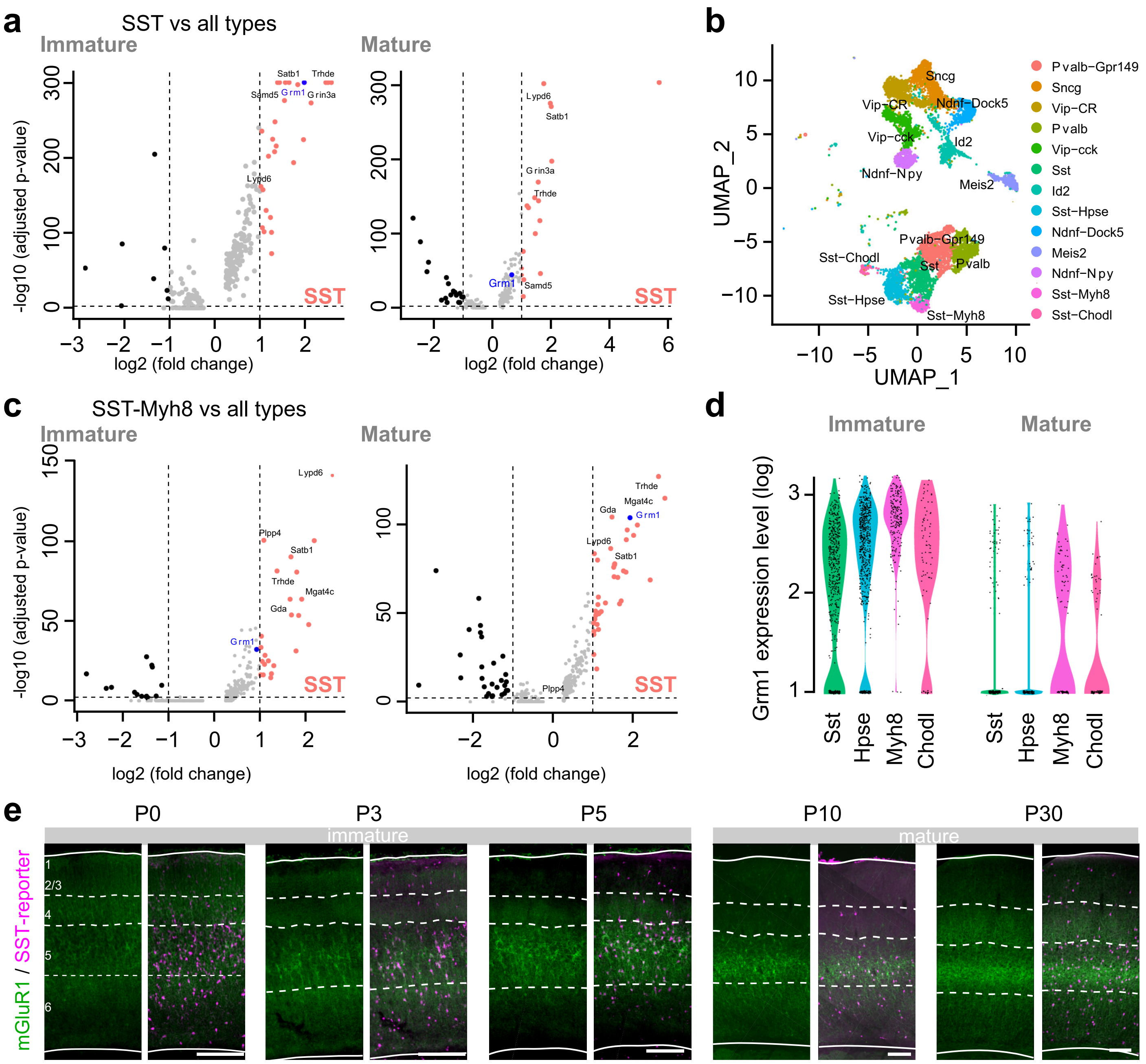


### Supplementary Fig. 3: mGluR1 expression in postnatal GABAergic interneuron

**a**, Differential gene expression of SST cIN cluster compared to every other major cIN class using Seurat non-parametric Wilcoxon rank sum test at P2 and P10. Other cIN populations were defined from other clusters defined in Fig. 3a, Grm1 is highly specific to SST cINs, more particularly at P2. **b**, UMAP representation of the integration of P2 and P10 cortical GABAergic neurons scRNA-seq public databases (Mayer et al., 2016 – P10; Allaway et al., 2019 – P2) allowing for the identification of SST cINs and other major cIN classes at both stages and expression markers described in Mayer et al. 2016 and Wu et al., Biorxiv 2022. **c**, Differential gene expression of the L5-specific SST-Myh8 cluster compared to every other cluster as described in (a) using Seurat non-parametric Wilcoxon rank sum test at P2 and P10. Compared to all other major clusters as defined in Fig. 3a, Grm1 is highly specific to L5-SST cINs, more particularly at P2. **d**, Violin plots representing the levels of Grm1 expression in all cells composing SST cIN clusters, showing that at P2, Grm1 is homogeneously expressed in all subtypes, while some specificity for deeper-layer SST cINs are visible at P10. **e**, Immunofluorescence of mGluR1 in S1 cortex. Expression is present as early as P0 and is high throughout development, with a soma to neurites redistribution of the receptor. Scale bars: 100 µm.

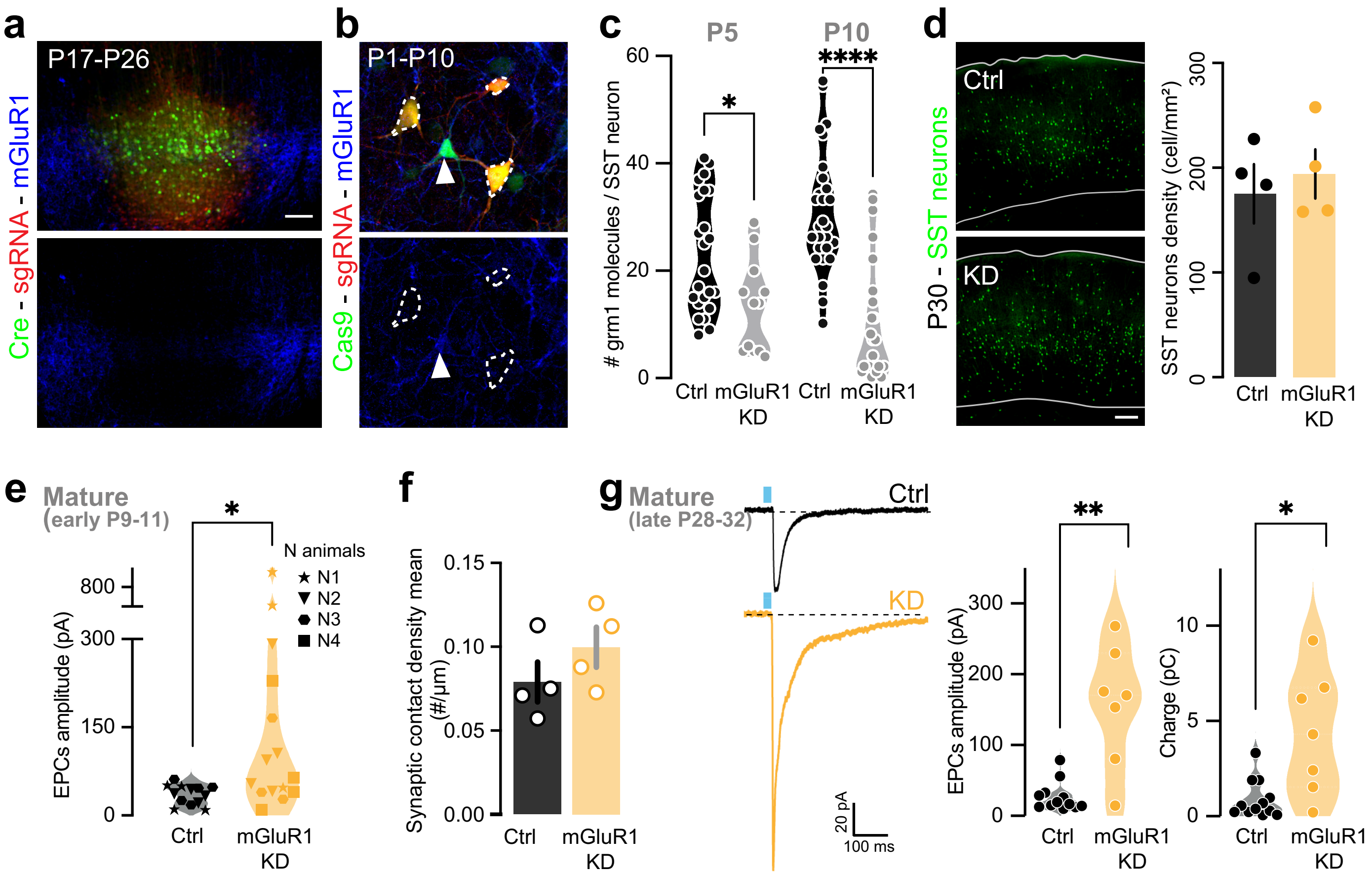


**Supplementary Fig. 4: CRISPR deletion of mGluR1 in SST cINs and TC input maturation**

**a**, gRNA validation: AAV-driven non-specific Cre-eGFP, AAV-driven sgRNA-DIO-dTomato and mGluR1 immunostaining reveal the deletion of mGluR1 protein expression after 10 days in the S1 injected region. Scale bar: 100 µm. **b**, Illustration of mGluR1 expression KD in cells infected by sgRNA (tdTomato). Cre-dependent Cas9+ cells without sgRNA do not exhibit mGluR1 deletion. **c**, Quantification of Grm1 RNA molecules per SST cINs using smFISH showing downregulation in the mGluR1 CRISPR KD cells at P5 and P10. Student t-tests (P5 p-value=0.0117, Ctrl: 23.09±2.29 n=23 N=2, KD: 13.33±2.48 n=12 N=2); (P10: p-value<0.0001, Ctrl: 30.89±2.17 n=27 N=2, KD: 10.35±2.35 n=23 N=2). **d**, Density of SST neurons/mm$^2$ of S1 showing no difference in number between Control and KD conditions at P30. Non-significant Student's t-test (Ctrl: 175.1±28.34 N=4; mGluR1 KD: 194.1±23.48 N=4). **e**, EPSC amplitudes from Fig. 4d, labeled for biological N (n cells recorded per animal). **f**, Average per animal of synaptic contact density onto SST cINs in control and mGluR1 KD SST cINs (Ctrl 0.079±0.012 N=4; mGluR1 KD 0.100±0.012 N=4). **g**, Averaged traces of example response from SST cIN responses to VB fiber stimulations at P30. EPSC peak amplitudes of control and mGluR1 CRISPR KD SST cINs in response to VB fiber stimulations at P30. Mann-Whitney test (p-value=0.0026; Ctrl from Supp. Fig. 1c: 26.92±6.01; mGluR1 KD 156±32.46). EPSC charges of control and mGluR1 CRISPR KD SST cINs in response to VB fiber stimulations at P30. Mann-Whitney test (p-value=0.0250; Ctrl: 1.05±0.33; mGluR1 KD 6.32±1.71).

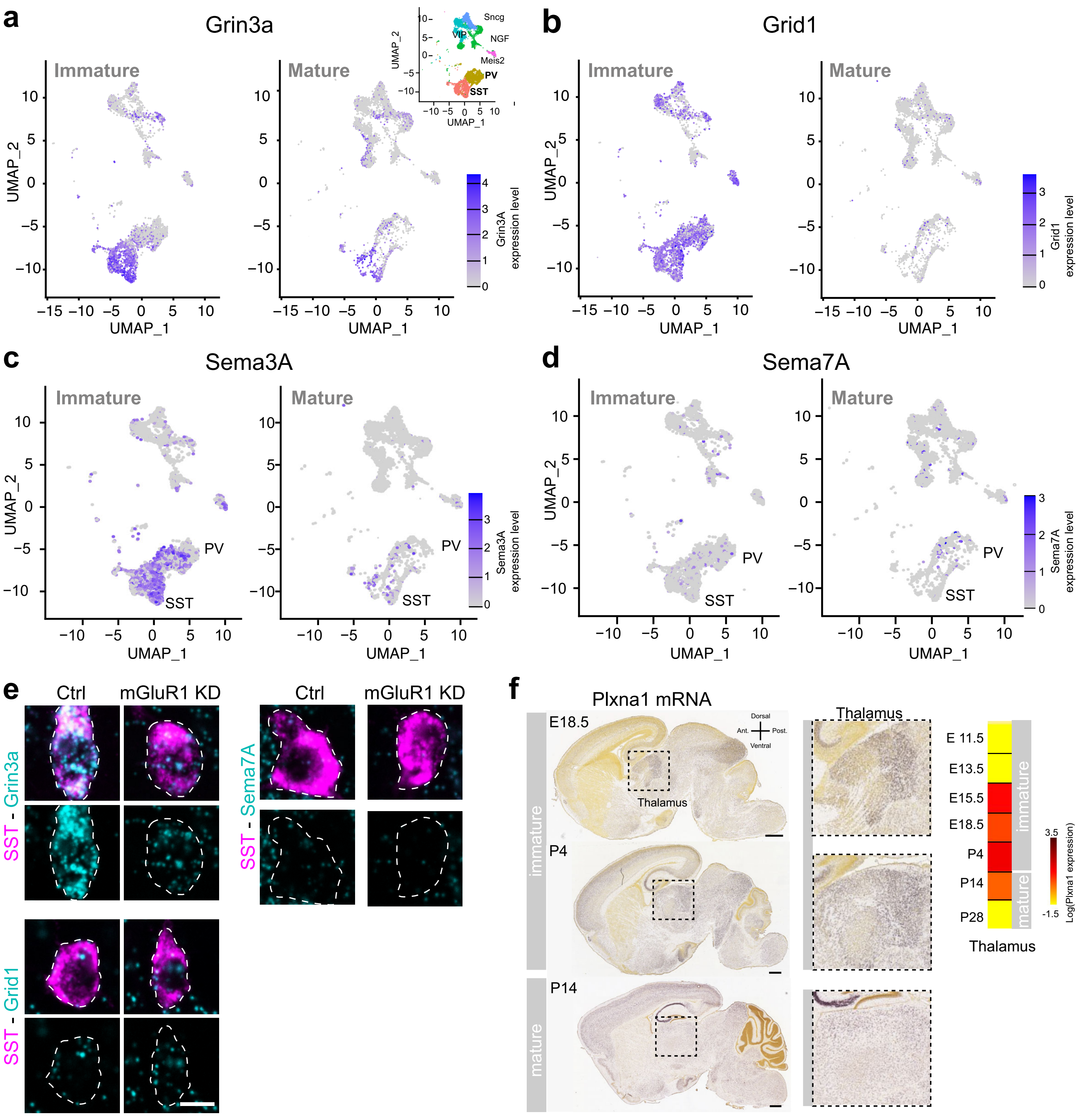


**Supplementary Fig. 5: Expression patterns of gene candidate for transient TC input regulation**
**a-d**,UMAP representation of P2 and P10 scRNAseq with expression levels of Grin3a, Grid1, Sema3A and Sema7A. **e**, Quantification of Grin3a, Grid1 and Sema7A transcripts per SST cINs, using smFISH and SST cINs labeled with Sst immunostaining at P10 in Control and mGluR1 KD. Scale bar: 10 µm. **f**, *In situ* hybridization on sagittal sections showing Plexin A1 expression in the thalamus at E18.5, P4 and with absence of expression at P14. Images and level of expression from the Allen Developing Mouse Brain Atlas. https://developingmouse.brain-map.org/.

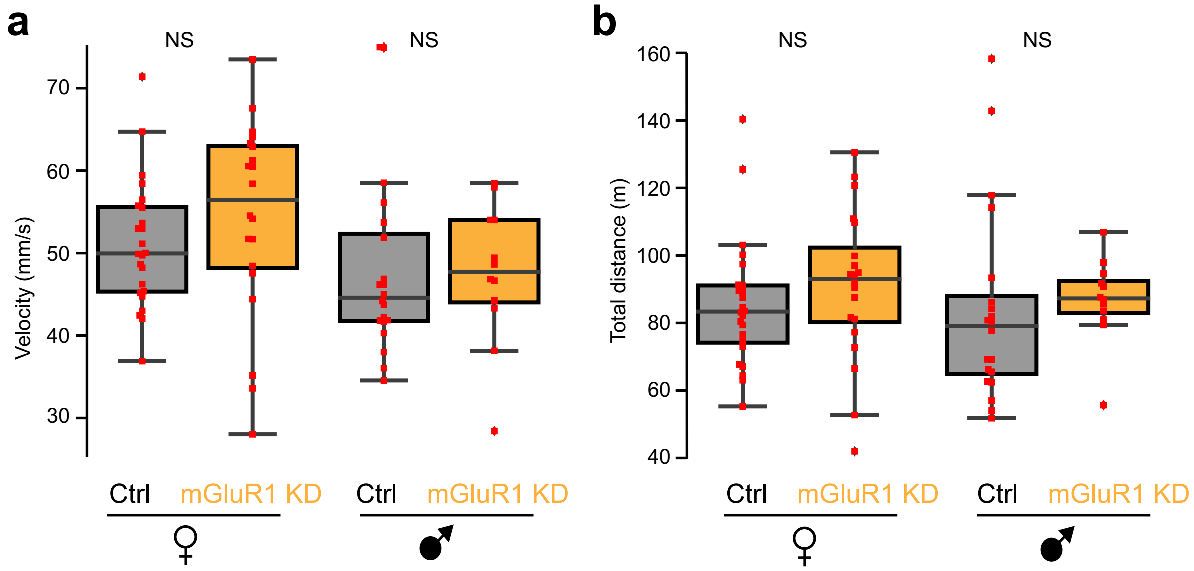


**Supplementary Fig. 6: Velocity and total distance traveled are not significantly different between mGluR1 KD and Control animals**

**a**, 2D velocity of individual mice in each sex, between mGluR1 KD and Control animals. Mann-Whitney U test was performed for 2D velocity between KD and Control in each sex. Female: test statistic=176, p=0.07; male: test statistic=99, p=0.21. **b**, Total distance traveled of individual mice in each sexbetween mGluR1 KD and Control animals. Mann-Whitney U test was performed for the total distance traveled between KD and Control in each sex. Female: test statistic=187, p=0.11; male=81, p =0.07.

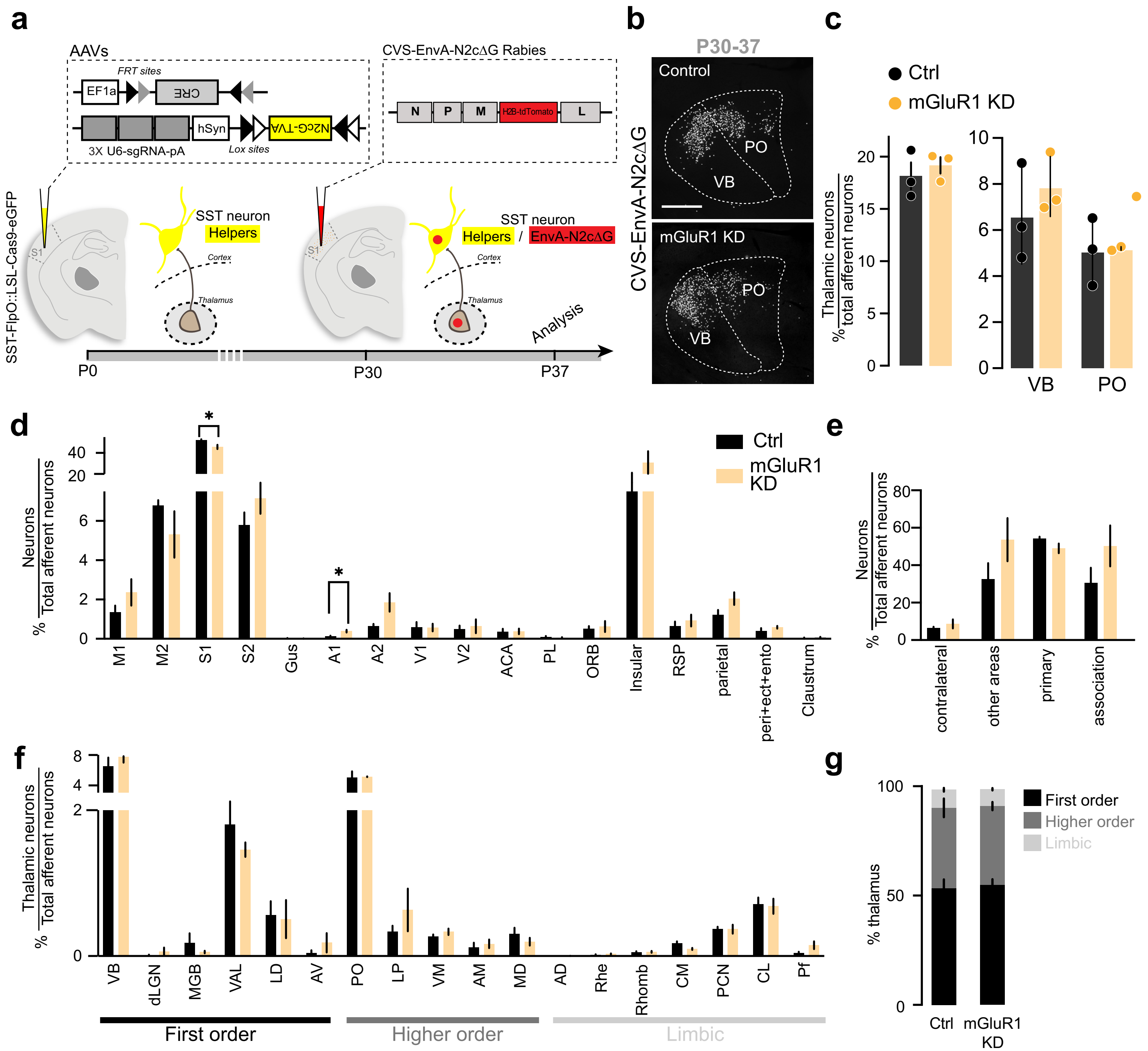


**Supplementary Fig. 7: Monosynaptic rabies retrograde labeling shows that removal of mGluR1 from SST cINs does not qualitatively affect their afferent connectivity**

**a**, Monosynaptic rabies tracing strategy. Cre-dependent rabies helpers (DIO-TVA-N2cG) are AAV-driven, together with mGluR1 CRISPR sgRNA. EnVA-pseudotyped CVS N2c rabies (RV) expressing the nuclear tdTomato were injected to infect SST cINs expressing RV-helpers. Experimental timeline: AAVs were injected at birth. RV were injected at P30 and brain collected for retrograde labeling quantification one week later. **b**, Nuclear tdTomato RV-retrograde labeling from SST cINs deleted for mGluR1 (KD) in the thalamus. Scale bar: 200 µm. **c**, The number of thalamic neurons normalized to the total of retrogradely labeled neurons from Control or KD SST cINs is not significantly different. Student t-test (Ctrl: 18.16±1.30 N=3; mGluR1 KD: 19.18±0.78 N=3). The proportion of VB and PO retrogradely labeled neurons is not significantly different either (Ctrl VB: 6.62±1.21; Ctrl PO: 5.09±0.85; mGluR1 KD VB: 7.89±0.75; mGluR1 KD PO: 5.20±0.04). **d**, Monosynaptic rabies retrograde labeling in cortical areas. The number of retrogradely labeled neurons in different cortical areas normalized to the total number of afferent neurons. Student's t-tests (p-value for S1=0.047; for A1=0.026).**e**, Same quantification as (d) with cortical areas grouped based on their hierarchy. Non-significant Student's t-tests. **f**, Same quantification as (c) from all thalamic nuclei. Non-significant t-tests. **g**, Proportion of retrogradely labeled neurons in the thalamus, based on their hierarchy: first order (FO) neurons, comprising VB neurons, higher order (HO) neurons, comprising PO neurons and limbic (Lb) neurons projecting to SST cINs is not significantly different between Control and KD SST cINs. Student's t-test (N=3).

| **Metrics** *(mean ± SEM)* | **DREADD_Gi** *(n = 12)* | **DREADD_Gq** *(n = 13)* | **GFP reporter (ctrl)** *(n = 14)* | **Kruskal-Wallis test p value** | **Dunn post-hoc test p value** |
|---|---|---|---|---|---|
| **Threshold (mV)** | -48.3 ± 1 | -50.7 ± 1.3 | -50.3 ± 2 | 0.025 | Gi *Vs* Gq : 0.036<br>Gi *Vs* Ctrl : 0.0086 |
| **IR (Mohm)** | 274 ± 32 | 397 ± 36 | 460.7 ± 46 | 0.017 | Gi *Vs* Gq : 0.023<br>Gi *Vs* Ctrl : 0.0076 |
| **Rheobase (pA)** | 22 ± 12.4 | 9 ± 6.3 | 16 ± 6.4 | 0.807 | - |
| **RMP (mV)** | -58.8 ± 2.5 | -59.2 ± 2.2 | -65 ± 2.5 | 0.499 | - |

**Supplementary Table 1.** Electrophysiological properties of SST neurons in DREADD-Gi, DREADD-Gq and GFP reporter control conditions. Related to Figure 2.

| | log2FC | adj. p value | | log2FC | adj. p value | | log2FC | adj. p value | | log2FC | adj. p value |
|---|---|---|---|---|---|---|---|---|---|---|---|
| Trhde | 2.6512 | 0 | Prkcg | 0.8845 | 1.5E-148 | Macrod2 | 0.8089 | 2.86E-94 | Il1rapl1 | -0.949 | 7.08E-76 |
| Grin3a | 2.4689 | 0 | Dlgap1 | -0.644 | 3E-148 | Grm5 | -0.829 | 5.49E-94 | Gm32647 | 1.1137 | 8.5E-76 |
| Grm1 | 2.1846 | 0 | Gm42418 | -0.844 | 4.2E-146 | Tnik | 0.6915 | 6.91E-94 | Fndc3b | 0.469 | 2.77E-75 |
| Grm7 | 1.7309 | 9E-264 | Cdh8 | 1.1399 | 3.7E-140 | Oprm1 | 0.7294 | 2.91E-93 | Rap1gap2 | 0.4884 | 4.76E-75 |
| Reln | 2.4522 | 4E-262 | Grin2a | 1.1563 | 1.1E-139 | Slc9a9 | 0.7228 | 3.57E-92 | Gm48321 | 0.6226 | 7.09E-75 |
| Mef2c | -1.421 | 8.1E-258 | Zeb2 | -0.82 | 2.3E-139 | Srrm4 | -1.082 | 3.99E-91 | Dync1i1 | 0.5714 | 7.39E-75 |
| Unc13c | 2.4189 | 3.4E-251 | Dlg2 | 0.7678 | 3.1E-139 | Rgs17 | 0.7214 | 5.74E-91 | Ildr2 | 0.4715 | 9.56E-75 |
| Ntm | 1.9046 | 1.3E-239 | Vcl | 0.8813 | 2.3E-138 | Ccser1 | 0.6672 | 1.32E-90 | Zfp536 | -1.122 | 2.43E-74 |
| Dscam | 1.8206 | 1.6E-238 | Gm48749 | 0.9381 | 1.8E-136 | Utrn | 0.7596 | 8.09E-90 | Gm15520 | 0.5067 | 5.1E-74 |
| Lingo2 | 2.2996 | 1.9E-237 | P3h2 | 1.037 | 3.8E-127 | Arhgap26 | 0.5983 | 5.36E-88 | Rasgef1b | 0.7396 | 6.91E-74 |
| Ralyl | 2.4288 | 1.2E-236 | Thsd7b | 1.127 | 3.1E-126 | Ank2 | 0.5147 | 5.47E-88 | Pld5 | 0.8966 | 1.25E-73 |
| Trpm3 | 2.1567 | 7.2E-236 | Slc24a2 | 0.9271 | 1.2E-125 | Slc2a13 | 0.6614 | 6.68E-88 | Man1a | 1.0014 | 1.33E-73 |
| Gm28175 | 1.7131 | 6.9E-234 | Ablim1 | 0.8501 | 2.6E-124 | Cnr1 | 0.5716 | 1.77E-87 | Gap43 | 0.4364 | 2.33E-73 |
| Erbb4 | -1.416 | 9.9E-218 | Lrtm1 | 0.8605 | 3.6E-123 | Trank1 | 0.5663 | 2.46E-87 | Pvt1 | 0.4949 | 8.89E-73 |
| Cacna2d3 | 1.5096 | 8.5E-213 | Npas1 | 0.7971 | 2E-122 | Grik3 | 0.7722 | 2.73E-86 | Tgfbr1 | 0.7181 | 9.08E-73 |
| Grik1 | 1.3547 | 1.8E-212 | Lhfpl3 | 1.0268 | 1.4E-116 | March1 | 0.762 | 3.76E-85 | Gm48727 | 0.4312 | 3.49E-72 |
| Ryr2 | 1.3414 | 1.2E-210 | Cttnbp2 | 0.8333 | 6.8E-115 | Erc2 | 0.5499 | 4.26E-85 | Far2 | 0.5022 | 4.52E-72 |
| Synpr | 1.6094 | 3E-207 | Nav2 | 0.9482 | 8.3E-115 | Otud7a | 0.6838 | 7.05E-85 | Pam | 0.6094 | 8.55E-72 |
| Frmpd4 | 1.6657 | 2.1E-201 | Mgat4c | 1.4924 | 1.7E-114 | Pde1a | 1.1113 | 8.13E-85 | Dlgap2 | -0.568 | 1.41E-71 |
| Cntn1 | 1.242 | 3.4E-200 | Elfn1 | 0.8547 | 5.8E-114 | Lsamp | 0.7401 | 1.57E-84 | Fam189a1 | 0.8666 | 1.88E-71 |
| Gm16083 | 1.1627 | 2.2E-194 | Tmtc2 | 1.3045 | 9.4E-113 | Lypd6 | 0.6965 | 5.97E-84 | Gabrg3 | -0.938 | 6.77E-71 |
| Nrxn1 | 0.7962 | 1.9E-184 | Tle4 | 0.807 | 1.8E-112 | Palm2 | 0.6079 | 1.35E-83 | Rps6ka5 | -1.078 | 8.02E-71 |
| Rbms1 | 1.3091 | 2.1E-183 | Thrb | 0.9663 | 7E-111 | Brinp1 | 0.7963 | 1.85E-83 | Csmd3 | 1.0723 | 1.22E-70 |
| Sh3rf3 | -1.252 | 1.1E-180 | Col19a1 | 1.0771 | 1.6E-109 | Rims1 | 0.5636 | 1.11E-82 | Nhs | 0.614 | 1.37E-70 |
| Lrfn5 | 1.4583 | 7.3E-179 | Sash1 | 0.7246 | 9.1E-106 | Pcdh17 | 0.9443 | 3.44E-82 | Rbfox1 | -0.603 | 1.67E-70 |
| Samd5 | 1.1551 | 1E-174 | Slc35f4 | 0.8099 | 5.3E-105 | Ncam2 | 1.1641 | 1.67E-81 | Gm48003 | 0.4881 | 2.52E-70 |
| Snhg11 | 1.323 | 1.8E-174 | Ctnnd2 | 0.5574 | 1.4E-104 | Gm10475 | 0.5254 | 2.31E-81 | Pou3f3 | 0.3979 | 2.68E-70 |
| Fam19a2 | -1.982 | 4.2E-174 | Grik2 | 0.7685 | 2.1E-103 | Csmd1 | 0.6681 | 4.09E-81 | Dnm3 | 0.545 | 2.86E-70 |
| Pfkp | 1.0734 | 2.9E-170 | Kcnb2 | 0.7705 | 8.8E-103 | Pclo | 0.6092 | 6.98E-81 | Tmeff2 | 0.9011 | 5.36E-70 |
| Prkg1 | 1.4584 | 1.8E-167 | Asic2 | 1.4546 | 1.5E-102 | Tox | 1.0163 | 1.11E-80 | Gm15155 | 0.7989 | 5.78E-70 |
| Satb1 | 1.2132 | 2E-166 | Gabrb1 | 0.9456 | 2.4E-102 | Lypd6b | 0.5383 | 8.67E-80 | Cobl | 0.4513 | 5.82E-70 |
| Meg3 | 0.8564 | 3.3E-163 | Kif26b | 0.8332 | 5.5E-102 | Plppr1 | 0.6653 | 9.04E-80 | Flrt2 | 0.7347 | 6.1E-70 |
| Scn9a | 0.9673 | 1.3E-162 | Slc4a10 | 0.6664 | 1.1E-101 | Lama4 | 0.582 | 1.01E-79 | Gm15398 | 0.6838 | 1.02E-69 |
| 4930473D10 | 1.1288 | 2E-161 | Nalcn | 0.7399 | 1.1E-101 | Csmd2 | 0.6342 | 1.53E-79 | Zbtb16 | 0.5592 | 2.57E-69 |
| Adcy2 | 1.3937 | 3.1E-160 | Ppp1r14c | 0.5661 | 1.2E-101 | Cdh9 | 1.2687 | 4.08E-79 | Cacng3 | 0.586 | 6.59E-69 |
| Gda | 1.0199 | 1.1E-159 | Rapgef4 | 0.7755 | 2.1E-101 | Slc24a3 | 0.7814 | 7.88E-79 | Iqsec1 | 0.5761 | 1.69E-68 |
| Nxph1 | -0.656 | 8E-159 | Gria4 | -0.807 | 1.6E-99 | Slc6a15 | 0.5802 | 2.97E-78 | Carmil1 | 0.5445 | 2.7E-68 |
| Neto1 | 1.2057 | 1.1E-157 | Tspan7 | 0.6612 | 1.98E-98 | Brinp3 | 1.2046 | 3.18E-78 | Tcerg1l | 0.5006 | 3.02E-68 |
| Spon1 | 1.2553 | 1.8E-157 | 4930567K20 | 0.5873 | 2.22E-98 | Rasgrf2 | 0.5686 | 4.21E-78 | Kcnh8 | 0.4901 | 9.15E-68 |
| Hs6st3 | 1.7267 | 4.8E-157 | Tcf4 | -0.494 | 3.24E-98 | Ccdc85a | 0.608 | 5.26E-78 | Gpc5 | 1.6682 | 2E-67 |
| Pdzd2 | 1.1667 | 2.4E-153 | Cacna1e | 0.8066 | 5.97E-98 | Cadm2 | 0.6807 | 5.99E-78 | Usp29 | 0.6233 | 5.84E-67 |
| Pantr1 | 1.2628 | 7.4E-152 | Galnt14 | 0.636 | 2.24E-97 | 2900055J20 | 0.4917 | 2.32E-77 | Klhl14 | 0.6738 | 9.13E-67 |
| Rgs6 | 1.4596 | 1.6E-151 | Ptprr | 0.721 | 8.36E-96 | 9130015G15 | 0.541 | 1.62E-76 | Myo16 | 0.5502 | 2.76E-66 |
| Mcc | 1.0328 | 3.4E-151 | Tenm4 | 0.7753 | 1.45E-95 | Shisa6 | 0.6965 | 3.15E-76 | Ptprn | 0.3892 | 2.88E-66 |
| Kctd16 | 1.4122 | 8.5E-150 | Astn2 | 0.8961 | 4.56E-95 | Kctd8 | 0.7776 | 3.86E-76 | Bcl11a | 0.6576 | 3.04E-66 |
| Egfem1 | 1.8942 | 1.1E-149 | Kcnd3 | 0.7897 | 5.48E-95 | Car10 | 1.1078 | 4.52E-76 | Sh3d19 | 0.4966 | 3.4E-66 |
| Grid2 | 1.0557 | 2.2E-149 | Zfp385b | 0.8392 | 1.69E-94 | Mef2a | -1.009 | 6.62E-76 | Kcnt2 | 0.6261 | 5.35E-66 |

**Supplementary Table 2**: Top SST neuron markers compared to PV neurons at P2

| | log2FC | adj. p value | | log2FC | adj. p value | | log2FC | adj. p value | | log2FC | adj. p value |
|---|---|---|---|---|---|---|---|---|---|---|---|
| Sst | 4.8793 | 1.2E-163 | Spon1 | 1.105 | 9.46E-26 | Chmp4b | 0.6627 | 1.75E-18 | Paip2 | 0.6104 | 2.19E-14 |
| Mef2c | -1.522 | 1.21E-87 | Zeb2 | -0.993 | 1.32E-25 | Dynll2 | 0.7069 | 2.31E-18 | Txndc17 | 0.5143 | 2.33E-14 |
| Synpr | 2.1358 | 1.23E-78 | mt-Nd5 | -0.909 | 1.36E-25 | Ywhah | 0.6985 | 3.11E-18 | Ensa | 0.5723 | 2.5E-14 |
| mt-Nd1 | -1.12 | 2.42E-73 | Cdkn1a | 0.9884 | 1.45E-25 | Schip1 | 0.6491 | 3.17E-18 | Nrxn1 | -0.766 | 2.59E-14 |
| Trhde | 1.861 | 1.43E-69 | Fam19a2 | -1.476 | 1.48E-25 | Lingo2 | 0.6874 | 4.04E-18 | Smim14 | 0.5847 | 2.91E-14 |
| mt-Cytb | -1.044 | 2.37E-65 | Map1lc3a | 0.735 | 3.02E-25 | Cpne4 | 0.5596 | 4.89E-18 | Raly | 0.4353 | 3.41E-14 |
| Grin3a | 1.6788 | 3.75E-63 | Reln | 1.3845 | 4.08E-25 | Rab27b | 0.8279 | 5.44E-18 | Nr2f2 | 1.1434 | 4.25E-14 |
| Fxyd6 | 1.2822 | 1.28E-61 | Swi5 | 0.8585 | 8.4E-25 | Grik3 | 0.8025 | 6.85E-18 | Cnih2 | 0.5183 | 5.02E-14 |
| mt-Nd4 | -0.976 | 6.65E-58 | Gabrg3 | -1.268 | 1.3E-24 | Ahi1 | 0.6893 | 7.1E-18 | Tac1 | -1.315 | 5.55E-14 |
| Erbb4 | -1.646 | 1.06E-54 | Ptprd | -1.274 | 1.72E-24 | Scn9a | 0.7627 | 9.4E-18 | Smim10l2a | 0.5337 | 6.37E-14 |
| AI593442 | -1.727 | 1.14E-54 | Mea1 | 0.7757 | 3.34E-24 | Gabarap | 0.7025 | 9.74E-18 | Epha5 | -0.838 | 7.17E-14 |
| Rab3b | 1.391 | 6.12E-53 | Crip2 | 0.8296 | 5.58E-24 | Klf5 | 0.6641 | 1.34E-17 | Mllt11 | 0.5041 | 8.92E-14 |
| Fxyd7 | 1.5698 | 3.23E-50 | Ypel3 | 0.8602 | 8.83E-24 | Nos1 | 1.7873 | 1.47E-17 | Cacng3 | 0.6462 | 9.48E-14 |
| Lypd6b | 1.3175 | 4.53E-49 | Dbpht2 | 0.908 | 1.17E-23 | Bex1 | 0.7668 | 1.53E-17 | Nedd8 | 0.5961 | 2.41E-13 |
| Gda | 1.2938 | 3.22E-48 | Elfn1 | 1.0011 | 1.48E-23 | Cplx2 | 0.8349 | 2.26E-17 | Lgals1 | 1.5429 | 2.71E-13 |
| Arhgdig | 1.0245 | 6.63E-48 | Rit2 | 0.6372 | 1.59E-23 | Ap2s1 | 0.7009 | 2.38E-17 | Pcp4 | 1.6772 | 2.72E-13 |
| mt-Nd2 | -0.949 | 3.6E-46 | Basp1 | 0.9242 | 4.11E-23 | Il1rapl1 | -1.442 | 2.45E-17 | Dscam | 0.556 | 2.78E-13 |
| Crhbp | 1.711 | 7.92E-44 | Gng3 | 0.8117 | 5.32E-23 | Sh3gl3 | 0.6337 | 3.26E-17 | Cstb | 0.4611 | 2.89E-13 |
| Cck | -2.111 | 1.11E-43 | Jund | 0.9044 | 7.01E-23 | Arpc5 | 0.6827 | 3.51E-17 | Bmyc | 0.5691 | 2.92E-13 |
| Lypd6 | 1.4584 | 5.19E-43 | Vsnl1 | 0.9234 | 9.81E-23 | Calm2 | 0.6901 | 4.3E-17 | Trps1 | -1.298 | 3.34E-13 |
| Slc6a1 | -0.964 | 2.59E-42 | Ppp1r14c | 0.7455 | 1.08E-22 | Dusp26 | 0.6816 | 4.36E-17 | Ndufa4 | -0.466 | 3.79E-13 |
| Ildr2 | 1.0692 | 6.73E-39 | Thsd7a | -1.625 | 1.72E-22 | Lpl | -1.521 | 4.44E-17 | Il34 | 0.6914 | 3.82E-13 |
| Gap43 | 1.2287 | 1.13E-37 | Dynlrb1 | 0.75 | 1.9E-22 | Nrsn1 | 0.7095 | 5.3E-17 | Sf3b5 | 0.4905 | 4.21E-13 |
| Grm5 | -1.048 | 2.6E-37 | Igfbp5 | 0.7834 | 2.02E-22 | Ntm | 0.6119 | 5.95E-17 | Grin2a | 0.5617 | 4.74E-13 |
| Rgs17 | 1.3538 | 1.03E-36 | Npas1 | 0.7562 | 3.48E-22 | Mien1 | 0.6662 | 1.21E-16 | Atp6v1f | 0.5486 | 5.42E-13 |
| Neto1 | 1.0787 | 1.52E-35 | Cnr1 | 0.8465 | 5.83E-22 | H1fx | 0.5389 | 1.24E-16 | Pcdh18 | 0.5145 | 5.96E-13 |
| Satb1 | 1.1011 | 7.94E-35 | Runx1t1 | 0.9472 | 1.38E-21 | Hsbp1 | 0.6099 | 1.67E-16 | Timm8b | 0.5962 | 6.5E-13 |
| Alcam | -1.164 | 2.42E-34 | Calm3 | 0.6945 | 1.76E-21 | Kif21a | 0.6548 | 2.93E-16 | Kcnh7 | -1.144 | 6.67E-13 |
| Pde1a | 1.2899 | 2.14E-33 | Sh3bgrl3 | 0.8222 | 3.1E-21 | Unc13c | 0.6437 | 3.22E-16 | Ppp1r11 | 0.5737 | 7.46E-13 |
| Rpp25 | 1.0664 | 7.82E-33 | Tmem91 | 0.7924 | 4.22E-21 | Itpr1 | -1.261 | 3.68E-16 | Hint1 | 0.5423 | 9.08E-13 |
| Atp1b1 | -0.74 | 1.75E-32 | Tspan7 | 0.7348 | 6.42E-21 | Dleu7 | 0.6229 | 3.87E-16 | Pmm1 | 0.4781 | 1.1E-12 |
| C1ql1 | -3.327 | 2.47E-32 | Elof1 | 0.6447 | 6.42E-21 | AI413582 | 0.5775 | 7.61E-16 | Cort | 1.2474 | 1.23E-12 |
| Pitpnc1 | 0.9478 | 5.45E-31 | Ctxn1 | 0.7738 | 9.46E-21 | Snrpn | 0.6307 | 7.69E-16 | Tcf4 | -0.501 | 1.4E-12 |
| 2900055J20 | 0.8992 | 6.57E-31 | Camk2n2 | 0.7951 | 2.17E-20 | Cdc42ep5 | 0.5672 | 9.37E-16 | Nsg1 | -0.531 | 2.73E-12 |
| Pou3f3 | 0.9323 | 3.26E-30 | Lrfn5 | 0.7023 | 3.1E-20 | Ache | 0.7531 | 1.38E-15 | Nfib | -1.279 | 2.87E-12 |
| Npy | 4.0553 | 1.13E-29 | Man1a | 0.7329 | 4.08E-20 | Shisa6 | 0.5908 | 1.49E-15 | Gtf2h5 | 0.5211 | 3.55E-12 |
| Ralyl | 0.8879 | 2.03E-29 | Fam46a | 0.7121 | 6.77E-20 | Srp9 | 0.5826 | 1.54E-15 | H1f0 | 0.5805 | 3.62E-12 |
| Nap1l5 | 1.0928 | 3.13E-29 | Grm1 | 0.7176 | 7.43E-20 | Polr2f | 0.5559 | 2.12E-15 | 2900011O08 | 0.6056 | 3.83E-12 |
| Bcl11a | 1.0064 | 4.66E-29 | Dynll1 | 0.7799 | 9.28E-20 | Pcdh19 | -1.174 | 2.13E-15 | Zcchc17 | 0.497 | 4.54E-12 |
| Camk2n1 | 1.1393 | 5.58E-29 | Vamp2 | 0.7395 | 1.02E-19 | Myl12b | 0.6362 | 2.77E-15 | Dtd1 | 0.471 | 5.45E-12 |
| Bex2 | 0.9478 | 1.5E-28 | Ap1s1 | 0.7153 | 1.34E-19 | Cystm1 | 0.6197 | 3.28E-15 | Cpne5 | 0.4096 | 6.34E-12 |
| 1500009L16 | 0.9026 | 2.03E-28 | Atp2b4 | 0.8356 | 1.45E-19 | B3gat2 | 0.5618 | 5.28E-15 | Cib2 | 0.591 | 1.03E-11 |
| S100a10 | 1.2202 | 3.47E-28 | Pfn2 | 0.7285 | 1.71E-19 | 1110008P14 | 0.6521 | 5.49E-15 | Shisa4 | 0.4746 | 1.1E-11 |
| Atxn7l3b | 0.8456 | 6.88E-28 | Pea15a | 0.7773 | 2.34E-19 | Mme | -1.132 | 6.01E-15 | Nrip1 | 0.5571 | 1.12E-11 |
| Ctxn2 | 1.0392 | 6.95E-28 | Cbarp | 0.6684 | 4.87E-19 | Coa3 | 0.6247 | 6.84E-15 | N4bp2l1 | 0.4252 | 1.14E-11 |
| Calm1 | 0.9855 | 1.44E-27 | Penk | 0.9533 | 6.66E-19 | Rcan1 | 0.494 | 7.03E-15 | Arhgef15 | 0.4584 | 1.2E-11 |
| Zwint | 0.9148 | 1.89E-27 | Chd3os | 0.7349 | 6.68E-19 | Rgs6 | 0.5743 | 7.58E-15 | Cmas | 0.4762 | 1.63E-11 |
| Vgf | 1.0184 | 2.85E-27 | Fkbp1b | 0.6864 | 9.44E-19 | Zcchc12 | 0.6095 | 1.05E-14 | Mrfap1 | 0.4865 | 1.75E-11 |
| Sstr1 | 0.9616 | 6.39E-27 | Nnat | 1.5594 | 1.63E-18 | Chodl | 1.2617 | 1.47E-14 | Tceal8 | 0.5158 | 1.9E-11 |
| Gng4 | 0.7954 | 1.36E-26 | Zrsr2 | 0.6701 | 1.69E-18 | Igf1 | 0.7177 | 1.65E-14 | … | … | … |
| Hap1 | 0.8979 | 4.01E-26 | Stxbp6 | 0.6186 | 1.73E-18 | Kcnmb4 | 0.4409 | 1.71E-14 | Sema3a | 0.3633 | 0.000124 |

**Supplementary Table 3:** Top SST neuron markers compared to PV neurons at P10

| **Metrics** *(mean ± SEM)* | **mGluR1 KD** *(n = 9)* | **Control*** *(n = 14)* | **Mann-Whitney U test p value** *(two-sided)* |
|---|---|---|---|
| **Threshold (mV)** | -43.1 ± 1.08 | -41.9 ± 1.5 | 0.175 |
| **IR (Mohm)** | 537 ± 64 | 548 ± 55 | 0.578 |
| **Rheobase (pA)** | 33.3 ± 6.2 | 28.9 ± 3.7 | 0.699 |
| **RMP (mV)** | -49.9 ± 2.1 | -48 ± 1.9 | 0.462 |

**Supplementary Table 4.** Electrophysiological properties of SST neurons in mGluR1 KD and control conditions. Related to Figure 4.

### FEMALE

| label | syllable | Ctrl Usage | KD Usage | KD-Ctrl delta | Ctrl Velocity | KD velocity |
|---|---|---|---|---|---|---|
| groom | 44 | 0.009 | 0.005 | -0.004 | 0.749 | 0.782 |
| scrunch | 15 | 0.024 | 0.022 | -0.002 | 1.116 | 1.118 |
| rear | 28 | 0.017 | 0.016 | -0.001 | 1.532 | 1.615 |
| rear | 45 | 0.009 | 0.007 | -0.001 | 0.614 | 0.61 |
| rear | 61 | 0.005 | 0.005 | 0 | 0.739 | 0.72 |
| rear | 42 | 0.008 | 0.008 | 0 | 0.609 | 0.599 |
| rear | 41 | 0.008 | 0.008 | 0 | 0.895 | 0.934 |
| groom | 36 | 0.012 | 0.012 | 0 | 1.718 | 1.736 |
| rear | 60 | 0.003 | 0.004 | 0 | 0.488 | 0.474 |
| rear | 64 | 0.006 | 0.006 | 0 | 0.733 | 0.709 |

### MALE

| label | syllable | Ctrl Usage | KD Usage | KD-Ctrl delta | Ctrl Velocity | KD velocity |
|---|---|---|---|---|---|---|
| rear | 41 | 0.013 | 0.008 | -0.005 | 0.828 | 0.914 |
| groom | 36 | 0.013 | 0.009 | -0.004 | 1.748 | 1.805 |
| groom | 44 | 0.01 | 0.008 | -0.002 | 0.743 | 0.757 |
| rear | 61 | 0.006 | 0.004 | -0.002 | 0.674 | 0.729 |
| rear | 64 | 0.004 | 0.003 | -0.001 | 0.719 | 0.743 |
| rear | 28 | 0.016 | 0.015 | -0.001 | 1.481 | 1.516 |
| rear | 42 | 0.012 | 0.012 | 0 | 0.574 | 0.564 |
| rear | 60 | 0.006 | 0.007 | 0 | 0.481 | 0.475 |
| rear | 45 | 0.008 | 0.01 | 0.001 | 0.594 | 0.604 |
| scrunch | 15 | 0.018 | 0.023 | 0.006 | 1.153 | 1.055 |

**Supplementary Table 5:**

Syllable labels, usages and velocity for the syllables with top 10 absolute LDA absolute model weight, related to Figure 6

**Supplementary Videos 1-10**

Videos of examples for syllables 15, 28, 36, 41, 42, 44, 45, 60, 61, 64, the top 10 syllables utilized in the linear discriminant analysis (LDA) related to Fig. 6. Class labels of the syllables: scrunch (15) , rear up (28, 36, 41, 45, 61), groom (42) , mid rear (44, 64), groom (60).
